# Spontaneous Traveling Cortical Waves Gate Perception in Awake Behaving Primates

**DOI:** 10.1101/811471

**Authors:** Zachary W. Davis, Lyle Muller, Julio-Martinez Trujillo, Terrence Sejnowski, John H. Reynolds

## Abstract

Perceptual sensitivity varies from moment to moment. One potential source of variability is spontaneous fluctuations in cortical activity that can travel as a wave. Spontaneous traveling waves have been reported during anesthesia, but questioned as to whether they are relevant to waking cortical function. Using newly developed analytic techniques, we find spontaneous waves of activity in extrastriate visual cortex of awake marmosets (*Callithrix jacchus*). In monkeys trained to detect faint visual targets, the timing and position of spontaneous traveling waves, *prior to target onset*, predict the magnitude of evoked activity and the likelihood of detection. In contrast, spatially disorganized fluctuations of neural activity are much less predictive. These results reveal an important role for spontaneous traveling waves in sensory processing through modulating neural and perceptual sensitivity.

**One Sentence Summary:** Fluctuations in cortical activity often travel as waves, shape incoming sensory information, and affect conscious perception.

## Main Text

Our perceptual experience can be highly variable. A faint stimulus presented at perceptual threshold may be detected in one instance but go undetected at another time. What occurs within the nervous system to account for this difference? Cortical neurons emit variable spike patterns in response to repeated presentations of an identical stimulus (1–3). This variability is due, in part, to ongoing spontaneous fluctuations in the local network state that regularly show periods of high or low excitability (4–6) and are reflected in the local field potential (LFP) (7, 8). Spontaneous fluctuations have been observed under anesthesia to propagate like waves in visual (9–12), auditory (13), and somatosensory (14) cortex. However there is controversy over whether waking spontaneous fluctuations travel as waves or, if they do, whether they contribute meaningfully to waking cortical function (15). Here, we report moment-by-moment fluctuations of neural activity recorded in behaving, non-human primates propagate as waves across the extrastriate middle temporal (MT) visual area. Critically, these waves are generated endogenously, and are thus distinct from previous reports of sensory- and behavior-evoked waves (12, 16–19). Further, we find that spontaneous waves strongly regulate visual perception. In particular, when the local population is in a depolarizing state during traveling waves, neuronal responses and perceptual sensitivity are both elevated in monkeys performing a challenging visual detection task.

We chronically implanted spatially distributed multi-electrode arrays (Utah Arrays, Blackrock Microsystems) into motion selective visual area MT of two marmosets, whose lissencephalic cortical structure allowed us to record simultaneously from the majority of the cortical area (Fig. 1A). We measured neuronal receptive fields from single- and multi-unit spiking activity in monkeys as they maintained fixation (Fig. S1). We also examined LFPs, which are driven by synaptic currents in the vicinity of the electrode, and reflect the excitability of the local network (4, 7, 8). From the perspective of a single electrode at one point in cortex, the raw LFP spontaneously fluctuates with broad spectral energy (20). However, when viewed simultaneously across a cortical area, the fluctuation peaks and troughs do not occur synchronously (Fig. 1B). Frequently, the peak of a fluctuation moves coherently across the cortex with the spatiotemporal profile of a traveling wave (Movie S1). As these peaks and troughs of the LFP can correspond to epochs of relatively low and high excitability in the local network, we will refer to peaks and troughs as hyperpolarizing and depolarizing states, respectively. We hypothesized that, when organized as waves, these states might gate the flow of spiking activity through cortical circuits depending on their alignment to the timing of afferent inputs (Fig. 1C).

**Fig. 1.**
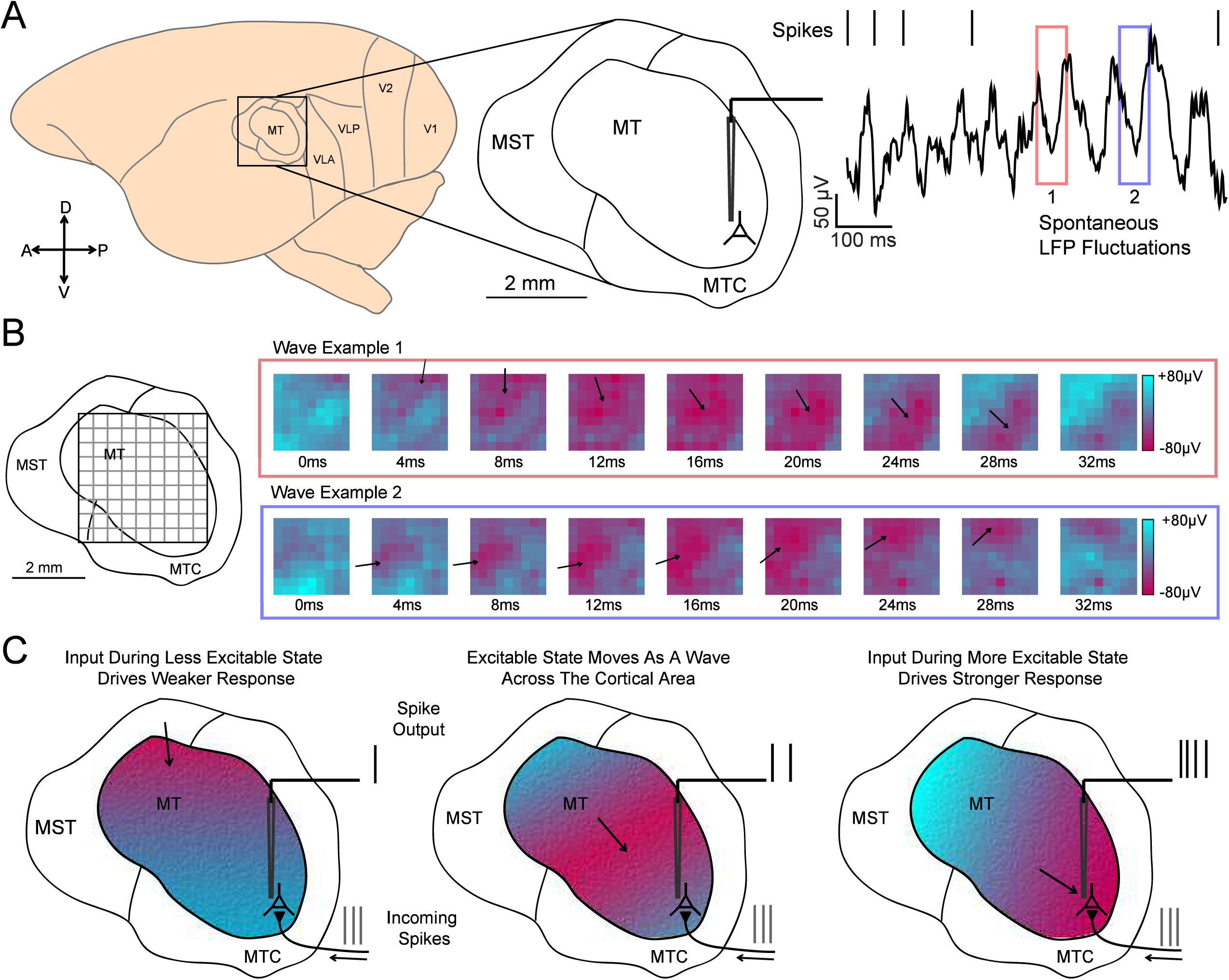
Spontaneous LFP fluctuations often travel as waves across the cortex. **(A)** Recordings were made from area MT of the common marmoset. The raw (1-100 Hz) local field potential (LFP) recorded from a single electrode is spectrally complex, with large, dynamic spontaneous fluctuations (red and blue boxes). **(B)** When viewed simultaneously across a multielectrode recording array, spontaneous fluctuations in the wideband (5-40 Hz) LFP often travel as a wave, showing spatiotemporal structure (example waves from red and blue boxes). The peaks and troughs of the wave correspond to spatially distinct regions of cortex that are more negatively (red) or positively (blue) potentiated. **(C)** Our mechanistic hypothesis: as waves traverse cortex, they alter excitability states, modulating the relative spiking output based on their alignment, with the depolarizing (red) and hyperpolarizing (blue) phases momentarily potentiating and, respectively, reducing sensitivity to incoming spikes.

Reliably detecting spontaneous waves in noisy multichannel data is challenging. Many wave detection techniques rely on spike-triggered averaging (12), spatial smoothing (10, 11), or narrowband temporal filtering (16, 19, 21) which can distort phase estimations of the underlying veridical fluctuation, giving false positives or unreliable measures of wave dynamics. Further, unlike during anesthesia, waking cortical dynamics are more complex, dominated by higher frequency, lower amplitude fluctuations that are more variable across the cortex (22, 23). To address this, we adapted a recently introduced statistical method for detecting traveling waves in noisy multichannel data (24) that is better suited to studying the dynamics of awake cortex (Fig. S2A). This method uses LFP phase to detect coherent flows of activity; however, while phase is conventionally analyzed only for narrowband oscillations, such as the theta (4-8 Hz) and alpha (8-13 Hz) frequency bands (25–27), the network fluctuations we observed were not stable, sustained narrowband oscillations (20) (Fig. 2B). Rather, they are broad in frequency content, and their frequency content shifts over space and time.

**Fig. 2.**
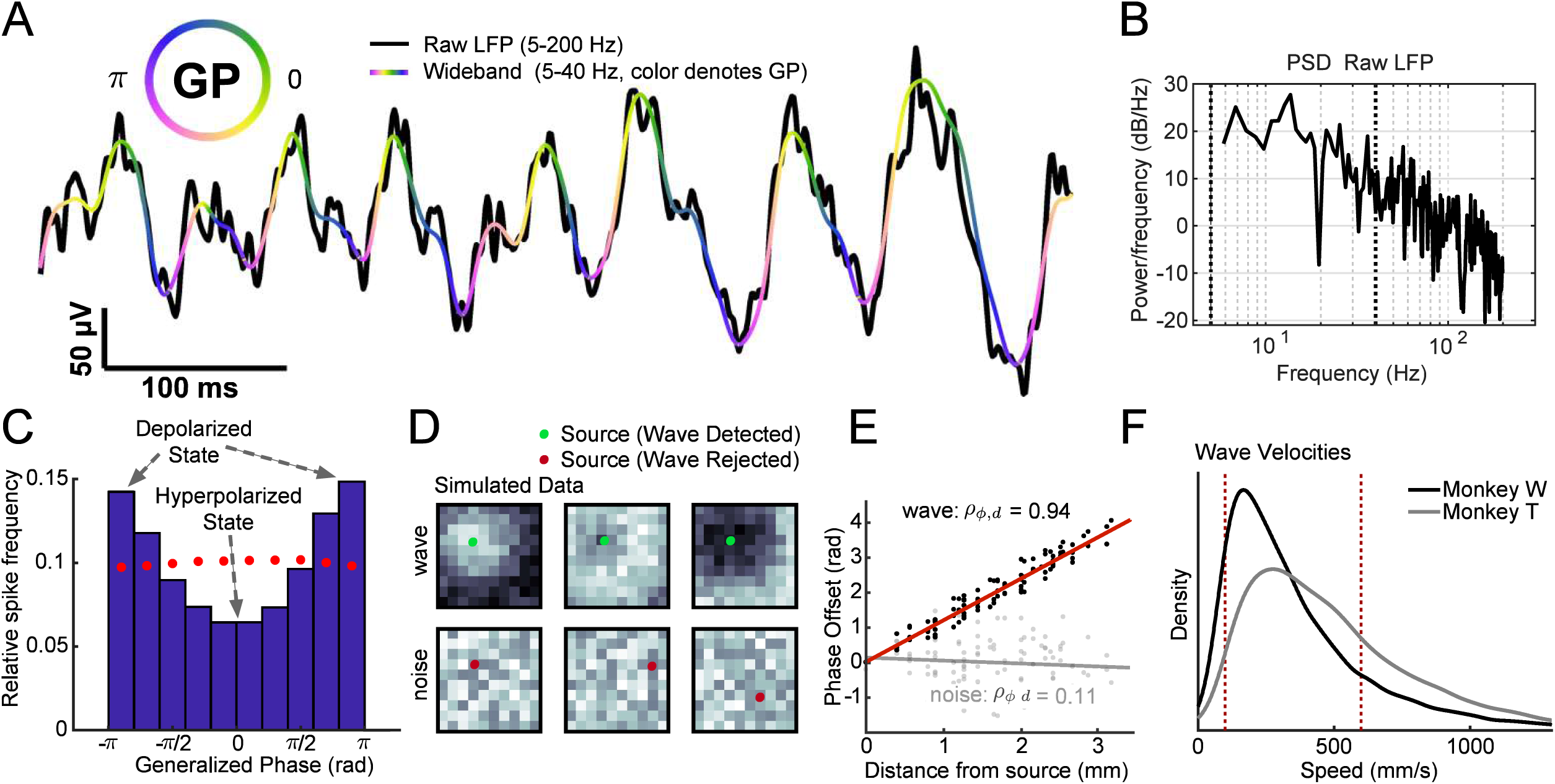
Spontaneous traveling waves modulate spontaneous spiking probability. **(A)** Illustration of the *generalized phase (GP)* measure used to detect spontaneous traveling waves. The GP (coded by color) describes the phase of the dominant fluctuation in the wideband (5-40 Hz, colored line) LFP at each moment in time, and is matched well to the raw LFP (5-200 Hz, black line). **(B)** Power spectral density for the raw LFP in (A), illustrating the broad spectral power in the signal. The bounds of the wideband filter are highlighted by the black dashed lines. **(C)** Spontaneous spikes are more likely to occur during depolarizing states (near ±π radians). Blue bars indicate the relative frequency of spikes occurring across GPs. Red dots indicate a control analysis: there is no phase relationship when the LFP is averaged across all electrodes, indicating the spatial specificity of spike-phase coupling. **(D)** A simulation demonstrating detection and rejection of traveling waves using our wave detection algorithm (see also fig. S2). Putative sources identified and tested by the algorithm are shown in green (detected candidate wave) and red (rejected candidate wave). **(E)** Waves are detected from the circular-linear correlation of phase offsets with distance from the putative source (black dots, linear fit in red line). The noise pattern shows no correlation with distance (gray dots, linear fit in gray line), and thus the wave is rejected. **(F)** Detected waves in both monkeys predominantly travel at speeds consistent with the conduction velocity of unmyelinated horizontal axons (100-600 mm/s, red dashed lines).

In order to track these fluctuating patterns, we expanded our wave detection method by developing a technique for computing the *Generalized Phase (GP)* of the wideband filtered (5-40 Hz) LFP (Fig. 2A). The wide frequency band captures the dominant fluctuating components of the LFP while excluding *1)* the lowest frequencies that are thought to reflect slow global changes such as arousal (23, 28) and *2)* higher frequencies so as to avoid contamination of the LFP by spike artifacts (29). Consistent with the identification of wave peaks and troughs as hyperpolarizing and depolarizing states, spontaneous spiking was strongly dependent on GP, with spike probability at the depolarizing state (±π radians) approximately twice that of the hyperpolarizing state (0 radians; p < 1 x 10^-5^, Rayleigh test for circular uniformity; Fig. 2B). The wideband filter captured the envelope of the LFP better than alpha and theta narrowband filters (Pearson’s correlation: wideband r = 0.91, significantly different from alpha r = 0.38 and theta r = 0.23, ɑ = 1 x 10^-5^ CI test; Fig. S3A) and spontaneous spiking activity was more strongly dependent on GP than on the phase computed using alpha or theta narrow-band filters (Fig. S3B, C). The strong spike-GP coupling was spatially specific (7, 30), as phases from adjacent electrodes were significantly less coupled to spike timing (monkey W: N = 23 sessions, 400 µM z = 4.70, p < 1 x 10^-5^, Wilcoxon signed-rank test; monkey T: N = 18 sessions, 633 µM z = 3.72, p < 1 x 10^-3^; Fig. S4).

Use of a wideband filter avoids phase distortions that could artificially produce waves or distort estimates of wave properties. Importantly, our wave detection algorithm was applied to spatially unsmoothed data, thus preserving as much of the veridical relationship between phase and spatial position as possible. Waves were either detected or rejected from the circular-linear correlation of phase with distance from putative sources on the recording array (Fig. 2C). Detected waves were then validated statistically against a randomly phase-shuffled null distribution of spatiotemporal patterns (>99th percentile, permutation test). Using this approach, we found spontaneous LFP fluctuations frequently travel as waves across the cortex while the monkey fixated a blank screen. It was not the case that larger fluctuations were more likely to be waves as there was no difference in the amplitude distribution between wave and non-wave fluctuations (Fig. S2D). Detected spontaneous waves traveled at speeds consistent with the conduction velocity of spikes traveling along unmyelinated horizontal projection axons (0.1-0.6 m/s; Fig. 2E) (31–33), suggesting waves propagate via the horizontal fibers that populate the superficial and deep layers of cortex. Further, though previous work found high contrast visual stimulation attenuates waves with a spike-triggered average (STA) analysis (12), our method found traveling waves during periods of fixation (−50 to +100 ms perisaccadic activity excluded) while the monkeys freely viewed high contrast naturalistic images (Fig. S5) indicating traveling waves are present during natural vision.

Do spontaneous waves play a role in perception? To test this, we trained two marmosets to perform a simple detection task. During the task, the monkeys were required to maintain fixation while awaiting the appearance of a low contrast (<2% Michelson contrast) 200ms drifting Gabor target that appeared at an unpredictable time at one of two equally eccentric locations, coinciding with retinotopic locations of channels on the array (Fig. 3A). Juice reward was provided if the monkey made a saccade to the location of a detected target within 500ms of target onset. For each monkey, target contrast was set to a value that was detected approximately 50 percent of the time (Fig. S6A).

**Fig. 3.**
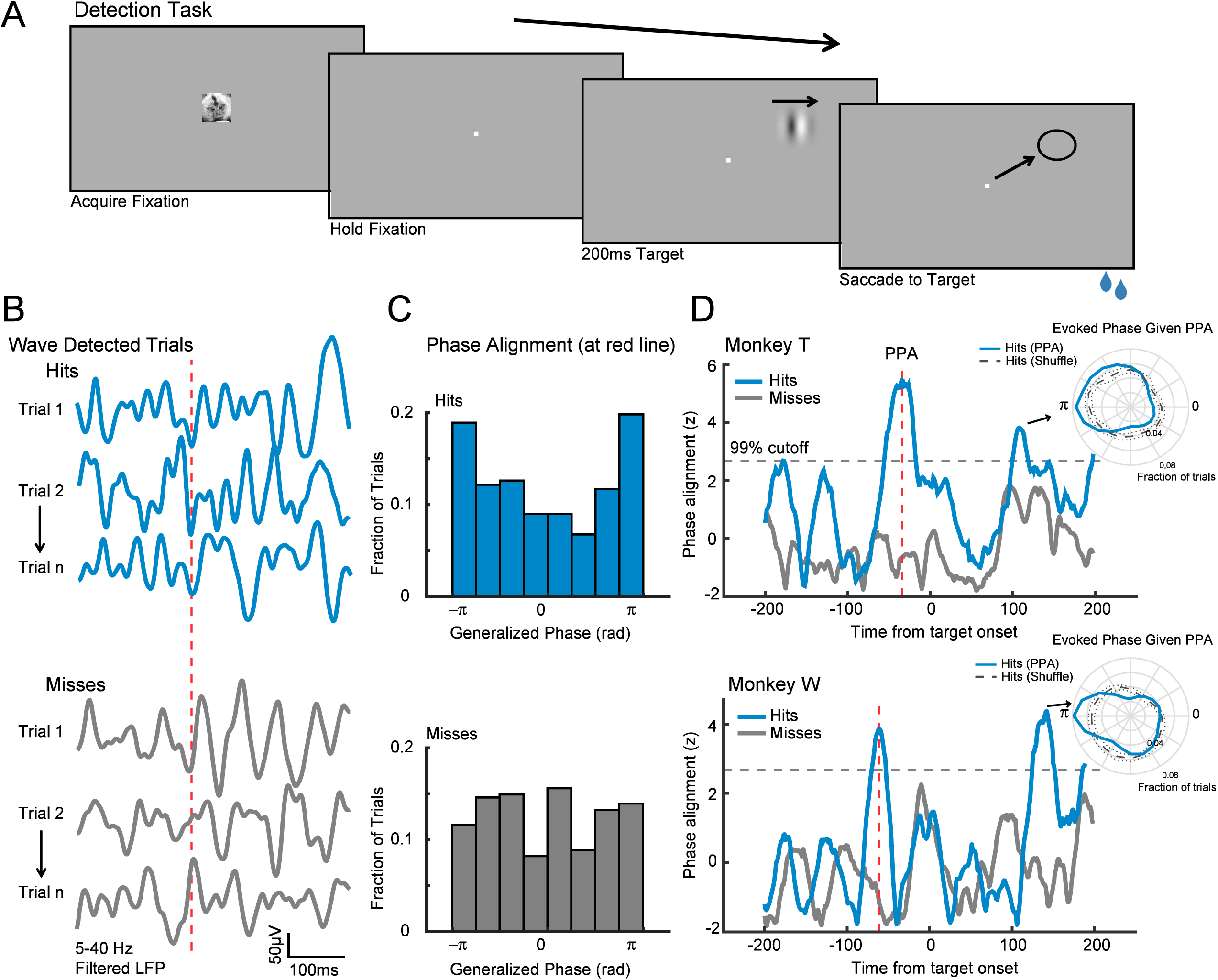
Waves facilitate detection when aligned with the retinotopic location of visual targets. **(A)** Target detection task. A marmoset face appeared, and when foveated it was replaced with a fixation point. Marmosets held fixation until a target (200ms drifting gabor) appeared at one of two possible locations after a random delay. The marmoset was rewarded with juice for making a saccade to the target location. **(B)** Example LFP traces from a target-aligned electrode (hits, blue; misses, gray) where waves were detected on the array at the time indicated by the red-dashed line. **(C)** Distributions of wave states across hit (blue, top) and miss trials (gray, bottom) at the time of the red dashed line in (B). Hit trials had a strong wave phase alignment across trials at this time. Miss trials had no phase alignment. Data in (B) and (C) are from monkey T. **(D)** The cross-trial wave phase alignment in (C) plotted for each moment in time around the onset of the target for monkey T (top) and monkey W (bottom). Each phase distribution was z-scored relative to shuffled phase distributions (gray dashed line: 99% cutoff from permutation control). The time of peak phase alignment (PPA, red dashed line) is shown for each monkey (−33 ms, monkey T; −60 ms, monkey W). **(D, insets)** Polar plots showing the distribution of phases during the evoked response of hit trials, conditionalized on having been at the aligned phase at PPA (blue), is more likely to be depolarized than expected from a shuffled distribution of hit trials (gray, dotted lines 95% CI).

We hypothesized that target detection should be facilitated when spontaneous waves align a depolarizing state with target locations. To test this, we collected trials on which waves were detected around the onset of a target and asked if detection probability varied with the state of the wave, measured at retinotopically aligned electrodes (Fig. 3B). We only included trials where fixation had been maintained for at least 300 ms prior to target onset to avoid including waves triggered by the saccade to fixation that initiated each trial (19). To avoid the confounding effects of waves generated by the appearance of the target, we only analyzed waves detected prior to the target-evoked response. There was a highly significant tendency for detected targets to have been preceded by a consistent wave state (Fig. 3C, D; peak −60 ms relative to target onset monkey W, p < 1 x 10^-3^, Rayleigh test; −33 ms monkey T, p < 1 x 10^-5^). We refer to this as the Pre-target Phase Alignment (PPA). Consistent with the mechanistic hypothesis in Figure 1C, the PPA wave state led to more consistently depolarized evoked responses, as trials close to the aligned phase at the time of PPA also exhibited more consistently depolarized LFP during the target-evoked response than expected from a distribution generated by shuffling all hit trials (permutation test, ɑ = 0.01, Fig. 3D insets). As the visual input was identical across the aligned and shuffled groups, the difference in evoked responses must be due to the prior wave state.

Consistent with findings that stronger target-evoked responses are correlated with better detection performance (34), we found that firing rates during the target-evoked response (70-200ms from target onset) were higher for hits than misses for both monkeys (monkey W: N = 17 single- and N = 56 multi-units; z = 3.36, p < 1 x 10^-3^, Wilcoxon signed-rank test; monkey T: N = 18 single and N = 49 multi-units; z = 4.44, p < 1 x 10^-5^; Fig. S6D, E). We hypothesized the depolarizing wave state predicted by the PPA modulates the magnitude of evoked activity, producing the stronger evoked responses correlated with improved detection. To test this, we calculated average multi-unit firing rates for the PPA that produced either depolarizing or hyperpolarizing states during the target-evoked response (Fig. 4A, B). Consistent with our hypothesis, the evoked firing rate was larger for waves that produced depolarizing states than hyperpolarizing states (p < 0.05 in both monkeys, one-tailed Wilcoxon rank-sum test). The phase at the time of PPA of similar amplitude fluctuations that did not meet our statistical criterion for waves (non wave) was not predictive and did not modulate the evoked firing rate (p > 0.25 in both monkeys). Thus, only the alignment of spontaneous traveling waves prior to target onset predicts a future depolarizing cortical state that increases the gain of the target-evoked response.

**Fig. 4.**
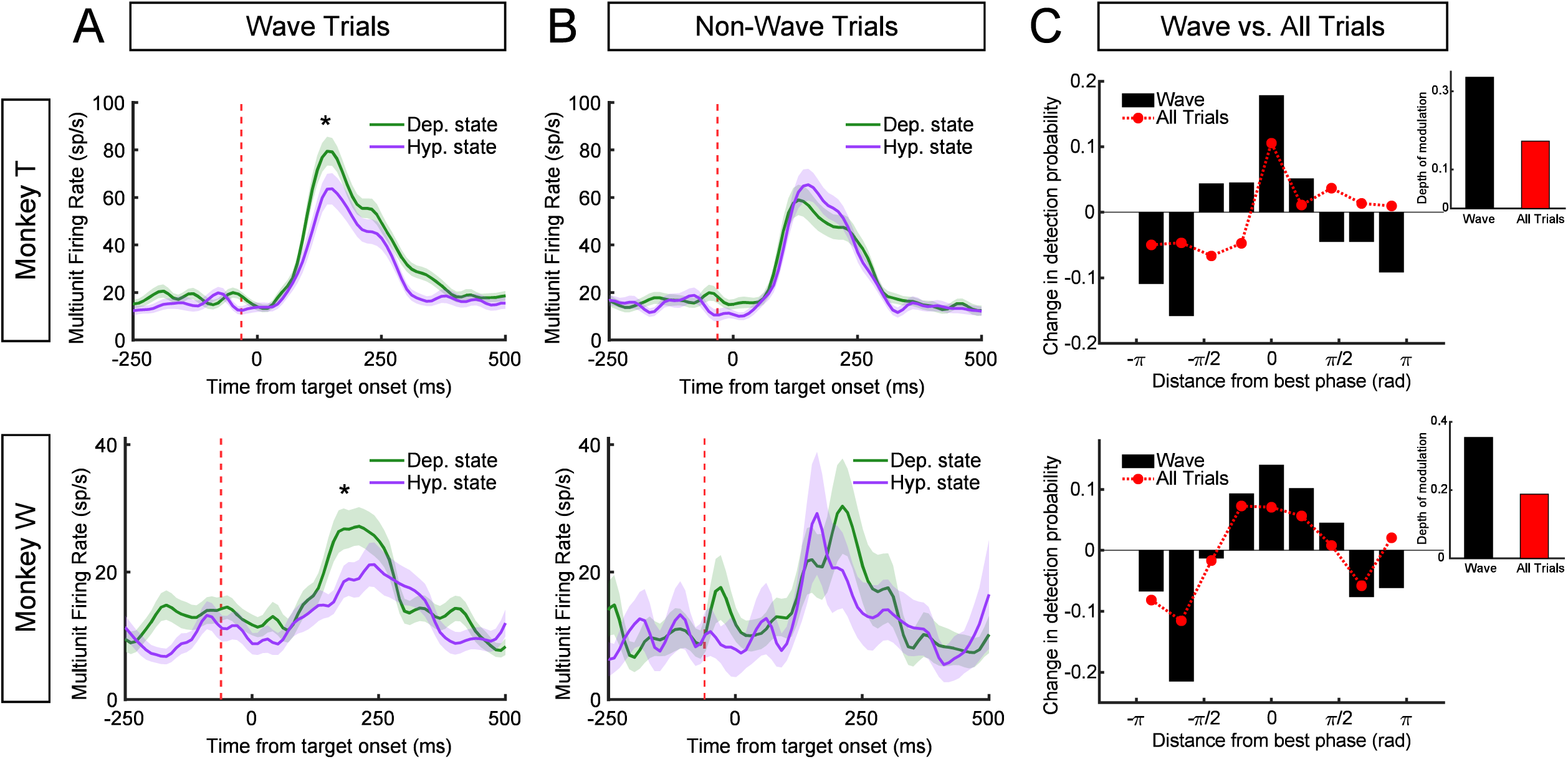
Wave state predicts target-evoked response magnitude and perceptual sensitivity. **(A)** Evoked multiunit responses are strongly modulated by wave state. Trials on which waves were detected at the time of PPA (dashed red line) were separated according to whether the wave state was depolarizing or hyperpolarizing. Average target-evoked responses following depolarizing wave states (green) are significantly greater than responses on trials following hyperpolarizing states (purple) for both monkey T (top row) and monkey W (bottom row; asterisks, p < 0.05). **(B)** Non-wave fluctuations did not modulate the magnitude of the evoked response. **(C)** Bar plots show the wave-state dependent change in detection probability centered on the best phase in each monkey. Red dots show weaker phase-dependent change in detection probability when all trials (wave, non-wave) are combined. The depth of modulation in detection probability given wave state is roughly double that for all trials (insets).

Finally, because the pre-target wave state predicts the magnitude of target-evoked spiking, we wished to quantify how well one could predict the monkey’s likelihood of detecting the faint target based solely on the knowledge of wave state at the time of PPA. We calculated the monkey’s conditional probability of target detection as a function of phase distance from the optimal aligned wave phase in each monkey (rotated to 0 radians). In both monkeys, the conditional probability of detection given wave state at the time of PPA reached a maximum for the optimal phases and a minimum for opposite phases (Fig. 4C), with a depth of performance modulation of 33% (monkey T) and 35% (monkey W) from peak to trough. If we take the perspective of a single electrode, with no knowledge about the state of the wave, but utilizing our more sensitive GP measure, there remains a strong phase dependant probability of detection, but significantly lower than when we exclusively look at trials that had predictive wave states before the target was presented (17% monkey T and 19% monkey W; p < 0.05 in both monkeys, confidence interval test).

Taken together, these results demonstrate that spontaneous traveling waves do occur in the neocortex of the awake monkey, they modulate sensory-evoked responses, and they gate perceptual sensitivity. The spontaneous traveling waves we detect are distinct from the large, slow-wave deflections reported during anesthesia or quiet wakefulness (35, 36). Rather, these waves are present during active vision, and their alignment preceding the presentation of the target predicts perceptual sensitivity to that target. Importantly, these wave effects are only apparent due to our measurement of the generalized phase, and could not be explained by latent narrowband oscillations embedded in the wideband signal. Narrowband filtering in alpha or beta bands fails to reveal any phase alignment predictive of perception (Fig. S7). The importance of waves to perception is further underscored by the fact that they are much more predictive of perceptual sensitivity than previous reports of pre-target alpha oscillation phase in visual detection (37–39) or theta oscillation phase in frontal-parietal networks during the deployment of attention (40–42). We speculate, given that we observe weaker predictive effects when we mix wave and non-wave trials, that the alpha and theta effects previously observed were in fact due to the undetected presence of traveling waves. This is supported by the recent discovery that alpha and theta oscillations travel as propagating waves across awake human cortex (21). If these two phenomena are related, this raises the intriguing possibility that the traveling waves we observe may also be coordinated across brain areas. Such coordination might explain how waves in MT are so strongly predictive of detection for stimuli that presumably activate other visual areas such as V1. These results have important implications for the neural organization of sensory processing demonstrating that, when viewed across the spatial extent of the cortex, fluctuations of cortical activity are neither purely synchronous nor spatially disorganized noise processes. Rather, neocortex exhibits propagating waves of activity that dynamically regulate neuronal responses and perceptual sensitivity.

## Acknowledgments

We would like to thank Michael Avery, Katie Williams, Sean Adams, and Mat LeBlanc for their contributions to this project. We would also like to thank Tony Movshon for his feedback in the early stages of this project.

## Author Contributions

Conceptualization, Z.W.D., L.M., J.H.R.; Data Curation, Z.W.D., L.M.; Formal Analysis, Z.W.D., L.M.; Funding Acquisition, Z.W.D., L.M., T.S., J.H.R.; Investigation, Z.W.D., L.M.; Methodology, Z.W.D., L.M., T.S., J.M.T., J.H.R.; Supervision, T.S., J.M.T., J.H.R.; Visualization, Z.W.D., L.M.,

Writing - original draft, Z.W.D., L.M., J.H.R.; Writing - review and editing, Z.W.D., L.M., T.S., J.M.T., J.H.R.

## Data and materials availability

An open-source code repository is available on BitBucket: http://bitbucket.org/lylemuller/wave-matlab. Reasonable requests for further access to data and materials will be accommodated.

## Materials and Methods

### Surgeries

One male (monkey W) and one female (monkey T) marmoset participated in this study. Each marmoset was fitted with a headpost for head stabilization and eye tracking. The headpost contained a hollow chamber housing an Omnetics connecter for a Utah array, which was chronically implanted in a subsequent surgery. For that surgery, a 7×10mm craniotomy was made over area MT (stereotaxic coordinates 2mm anterior, 12mm dorsal). An 8×8 (64 channel, monkey W) and 9×9 with alternating channels removed (40 channel, monkey T) Utah array was chronically implanted over area MT using a pneumatic inserter wand. The electrodes spacing was 400μM with a pitch depth of 1.5mm. The craniotomy was closed with Duraseal (Integra Life Sciences, monkey W) or Duragen (Integra Life Sciences, monkey T), and covered with a titanium mesh embedded in dental acrylic. All surgical procedures were performed with the monkeys under general anesthesia in an aseptic environment in compliance with NIH guidelines. All experimental methods were approved by the Institutional Animal Care and Use Committee (IACUC) of the Salk Institute for Biological Studies and conformed with NIH guidelines.

### Data Acquisition

Marmosets were trained to enter a custom-built marmoset chair that was placed inside a faraday box with an LCD monitor (ASUS VG248QE) at a distance of 48cm. The monitor was set to a refresh rate of 100Hz and gamma corrected with a mean gray luminance of 75 candelas/m^2^. Electrode voltages were recorded from the Utah arrays using two Intan RHD2132 amplifiers connected to an Intan RHD2000 USB interface board. Data was sampled at 30 kHz from all channels. The marmosets were headfixed by a headpost for all recordings. Eye position was measured with an IScan CCD infrared camera sampling eye position at 500 Hz. Stimulus presentation and behavioral control was managed through MonkeyLogic in Matlab. Digital and analog signals were coordinated through National Instrument DAQ cards (NI PCI6621) and BNC breakout boxes (NI BNC2090A). Neural data was broken into two streams for offline processing of spikes (single-unit and multi-unit activity) and LFPs. Spike data was high-pass filtered at 500 Hz and candidate spike waveforms were defined as exceeding 4 standard deviations of a sliding 1 second window of ongoing voltage fluctuations. Artifacts were rejected if appearing synchronously (within 0.5 ms) on over a quarter of all recorded channels. Segments of data (1.5 ms) around the time of candidate spikes were selected for spike sorting using principal component analysis through the open source spike sorting software MClust in Matlab (A. David Redish, University of Minnesota). Sorted units were classified as single- or multi-units and single units were validated through the presence of a clear refractory period in the autocorrelogram. LFP data was low-pass filtered at 300 Hz and down-sampled to 1000 Hz.

### Receptive Field Mapping

Receptive fields were mapped through reverse correlation. The marmoset was trained to hold fixation on an image (marmoset face, 1 degree diameter) presented to the center of the LCD monitor. A drifting Gabor (2 degrees diameter, spatial frequency: 0.5 cycles per degree, temporal frequency 10 cycles per second) was presented at a random position on the monitor between 0-18 degrees in azimuth and −15 to 15 degrees in elevation, drifting in one of 8 possible directions for 200ms, after which it disappeared. After a random delay drawn from an exponential distribution (mean 40ms), a new probe appeared and the pattern repeated until the marmoset broke fixation (defined as an excursion of 1.5 degrees from fixation) or viewed 16 probes. The marmoset was given a juice reward proportional to the number of probes presented. The receptive field for each unit recorded on the array was estimated by calculating the spike-triggered average (STA) stimulus that evoked the maximal response:

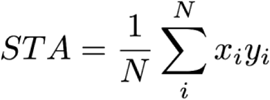

The STA is the sum of probe location x_i_ weighted by the spike count y_i_ within the time bin 40 to 200ms after probe onset, normalized by the number of all recorded spikes *N*. From the location of estimated receptive fields on each spiking channel, and the known topography of area MT in the marmoset (43), we estimated the relative position of each recording array in marmoset cortex (Fig. 1a). We excluded from the analysis the upper half of monkey W’s array as the recordings did not appear to be in area MT.

### Target Detection Task

Marmosets initiated each trial by fixating a marmoset face (a stimulus that naturally attracts marmoset gaze) that, upon fixation, transformed into a fixation point (0.15 dva). They were trained to hold fixation on the fixation point (1.5 degree tolerance) awaiting the appearance of a drifting gabor (4 degrees diameter; spatial frequency, 0.5 cycles per degree; temporal frequency, 10 cycles per second, drifting in one of up to 8 possible directions). After establishing fixation, the marmoset was required to hold fixation for a minimum duration (400 ms monkey W, 300 ms monkey T) to avoid contamination from waves caused by the saccade to the fixation point (19). The target only appeared if fixation was held for an additional random duration drawn from an exponential distribution (mean 200ms) to generate a flat hazard function. The target could appear at one of two locations selected based on receptive field mapping at equal eccentricity (7 degrees monkey W, 8 degrees monkey T). The target was presented for 200ms, after which the monkey had 300ms (for a total of 500ms) to saccade to within 2.5 degrees of the target center for a juice reward. On 10 percent of trials no target was presented, and the marmoset was rewarded for holding fixation to the trial end. The trial was classified as a miss if the marmoset broke fixation to a non-target location after the target had appeared, or if the marmoset held fixation until after the response window closed. If a saccade reached the target in less than 100 ms from target onset, the trial was rejected from analysis. Only trials from the preferred directions of motion for each unit were analyzed for that unit. Target contrast was selected as the value the marmoset correctly detected on average 50 percent of the time (mean 1.4 percent Michelson contrast for both monkeys). High contrast (10%) targets were presented on 10 percent of trials. If performance for these targets was below 70 percent, the session was rejected from analysis.

### Free Viewing Natural Images

Marmosets were headfixed and their gaze was monitored as in the previous tasks. Grayscale versions of naturalistic images were randomly interleaved and presented to the monkey. The monkey was free to look at the images, and after 10 seconds was given a juice reward. Saccades were identified from the velocity of the vertical and horizontal eye traces, and spontaneous fixations defined as periods where the eye velocity was 0. To detect waves we only included spontaneous fixation data from after 100 ms following the end of a saccade, and 50 ms before the start of a new saccade.

### Generalized Phase

The analytic signal paradigm was originally developed by Denis Gabor in 1946 (44), defining the concept of “instantaneous frequency” and “instantaneous phase” for non-stationary signals; however, due to several technical limitations, the analytic signal representation is commonly used strictly in the context of signals pre-treated with a tight narrowband filter (45). Here, we sought to address the technical limitations in the analytic signal to generalize its application beyond signals where tight narrowband filtering is appropriate. For this reason, we call our updated approach for non-stationary, wideband signals *generalized phase (GP)*.

Consider a real-valued signal *x_n_* ∈ **ℝ** for *n ∈ [1,2,…,N_s_]*, where *N_s_* is the number of samples in one recorded trial obtained at a sampling frequency *F_s_*. Given *x_n_*, its analytic signal representation (*X_n_*) is:

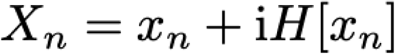

where *i* is the imaginary unit and *H[y_n_]* is the Hilbert transform (HT) of the signal *y_n_*. This representation can be obtained by implementing the HT operator as an FIR filter in time domain (46), or by using a single-sided Fourier transform approach (47, 48). Sinusoidal cycles appear in this representation as circular contours in the complex plane, while non-sinusoidal fluctuations appear as closed, quasi-circular contours. In this complex plane representation, phase is calculated numerically by the four-quadrant arctangent function.

The technical limitations in the analytic signal framework occur for two principal reasons. First, low-frequency intrusions effectively shift the representation by a complex constant, which has the critical effect of highly distorting phase angles estimated by the arctangent. As an initial step in the GP representation, then, we filter the signal within a wide bandpass (5-40 Hz), excluding low-frequency content. Note that this important step is distinct from narrow bandpass filtering (e.g. 8-13 Hz), as this approach preserves a significant portion of the signal spectrum, thereby minimizing waveform distortion and potential artifacts due to narrowband filtering of broadband noise (Fig. 2A and Supplementary Figure S3A). We then use the single-sided Fourier transform approach (47, 48) on the wideband signal and compute phase derivatives as finite differences, which are calculated by multiplications in the complex plane (18, 24, 49). Second, high-frequency intrusions appear in the analytic signal representation as complex riding cycles (49), which manifest as periods of negative frequencies in the analytic signal representation. As a secondary step in the GP representation, then, we numerically detect these complex riding cycles -- namely, *N_c_* points of negative frequency in the phase sequence *Arg[X_n_]* -- and utilize shape-preserving piecewise cubic interpolation on the next *2N_c_* points of *Arg[X_n_]* following the detected negative frequency epoch. The resulting representation captures the phase of the largest fluctuation on the recording electrode at any moment in time (Fig. 2A), without the distortions due to the large, low-frequency intrusions or the smaller, high-frequency intrusions characteristic of the *1/f*-type fluctuations in cortical LFP (50–52).

The GP represents a coherent numerical approach to the original analytic signal framework of Denis Gabor (44), suitable for implementation in modern digital signal processing applications. A complete description of this method, along with discrete analytical formulas for the GP representation, will be the subject of an upcoming work.

### Wave Detection

We employed a recently introduced statistical approach to detect spontaneous traveling waves in noisy multichannel recordings (18, 24), adapted to utilize GP. The advantage of GP is to capture the dominant fluctuation on each electrode at each point in time; further, it does not distort the signal waveform, as would occur with a narrowband filter. When these fluctuations are shared across electrodes and exhibit consistent phase offsets, the algorithm detects these patterns as traveling waves, as described below and illustrated in Figure S2A.

The wave detection technique occurs in three steps. First, the algorithm finds the time point nearest to each positive LFP peak on the array. This defines a flexible window in which we test for a spontaneous wave, where the phases are valid in a neighborhood around that time point. Secondly, the algorithm finds the most likely starting point for the wave, by finding the point that maximizes the divergence of the phase gradient in a smoothed version of the scalar phase field. This captures the point from which neural activity flows outward at each moment. Thirdly, with the putative source point found, the algorithm then quantifies how clearly activity is organized about this point, by calculating the circular-linear correlation with distance (ρ_φ,d_ ∈ [-1,1] (53)) across the whole electrode array, consistent with our observation that the wavelengths were long relative to the spatial extent of the array. Importantly, this step is done on the tested scalar phase field without spatial smoothing, which prevents smoothing artifacts from contaminating the results. Finally, a null distribution was constructed for ρ_φ,d_ by randomly shuffling phase values on the electrode array. Unless stated otherwise, a scalar threshold of 0.3 was used to detect waves throughout, which represented a conservative threshold on all constructed null distributions. For the analysis in Fig. 4C, we used the medians of the ρ_φ,d_ distribution in each monkey for the wave and non-wave states, respectively.

### Cross-Trial Phase Alignment

To quantify alignment of the GP across trials, we utilized the standard formulation for the Kuramoto order parameter:

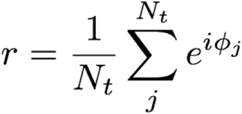

where *N_t_* is the number of trials in each condition (hit or miss), *i* is the complex unit, and φ_j_ is the GP at the tested time point. The order parameter ranges between 0 (uniform distribution of phase values) and 1 (identical phase values for each trial). To compare meaningfully between two sets of observations (hit and miss) with slightly different number of trials while accounting for the expected mean and variances of the order parameter at finite scales (54), all phase alignment values were put in z-score units of a null distribution computed from 10,000 iterations of the value from randomly selected trials, with the same number of observations.

### Conditional probability estimate

In order to understand how waves modulate the probability of target detection, we calculated the conditional probability of detection at each phase:

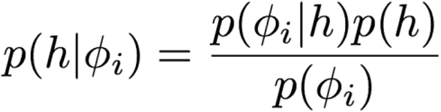

where h ∈ {0,1} is an indicator variable for target detection and Φ_i_ represents GP in bin i ∈ [1,N_b_], where N_b_ is the number of bins (nine throughout). To balance trial counts between wave and non-wave conditions, in this analysis we used the median of the ρ_φ,d_ distributions as the wave detection threshold. We then fit a sinusoid by least-squares estimate to the binned conditional probabilities in wave and non-wave states. Finally, significance of the difference in modulation amplitude between the two states was assessed in each monkey by comparing confidence intervals for each fit at the α = 0.05 level

## Supplementary Figures

**Fig. S1.**
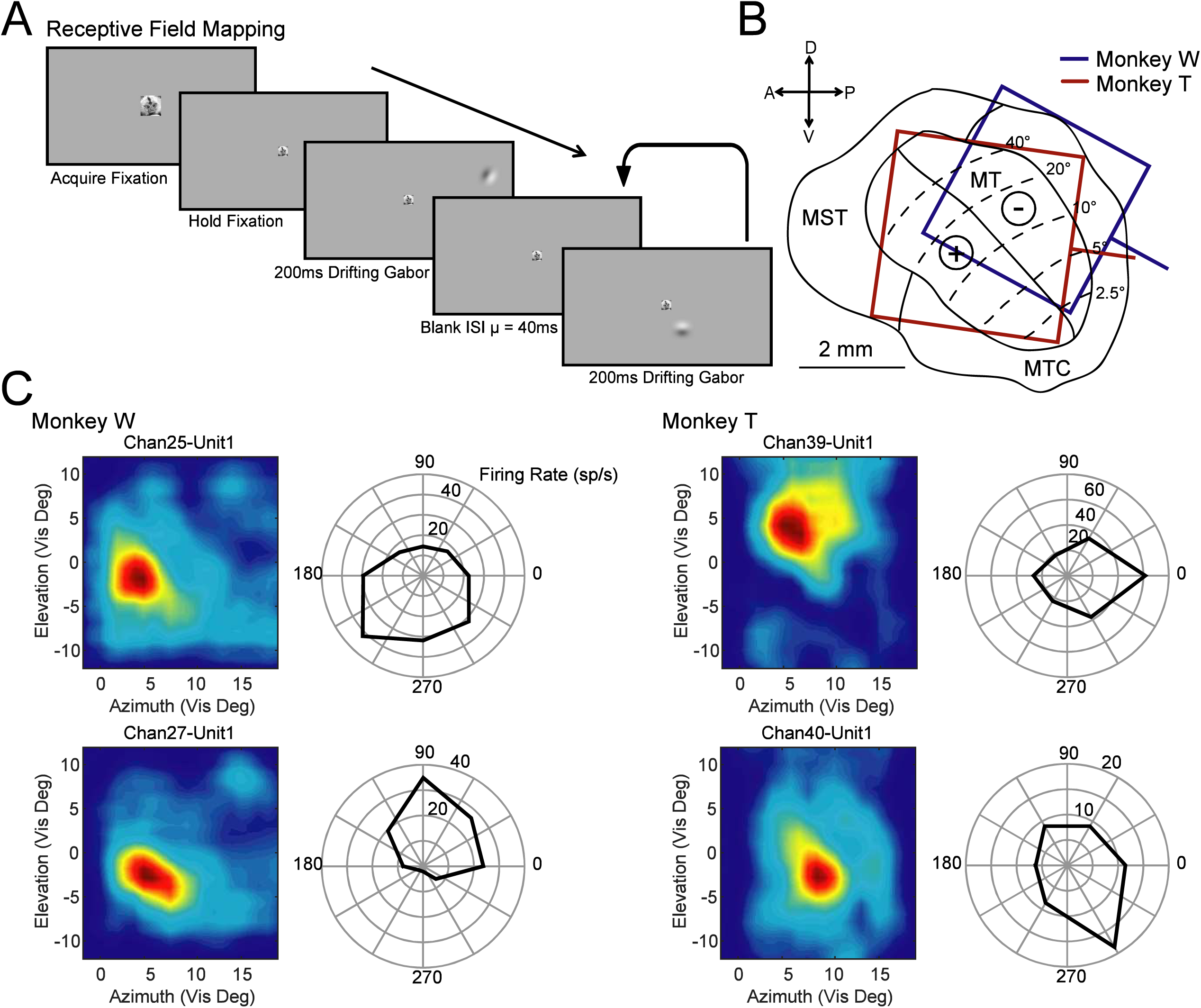
Retinotopic mapping and motion direction tuning is consistent with the anatomical organization and tuning preferences of marmoset MT. **(A)** Receptive fields for recorded units were measured by reverse correlation. Monkeys held fixation on a marmoset face while visual probes (drifting Gabors) appeared at random locations in the visual hemifield contralateral to the recording array. Each probe would appear, drift for 200ms, and disappear after which a new probe would appear in a new random location and the process would repeat until the monkey broke fixation. **(B)** The estimated position and orientation of Utah arrays in area MT based on retinotopy and histological examination for monkey W (blue) and monkey T (red). **(C)** Example receptive fields and their preference for motion direction are consistent with previous reports of marmoset MT (43).

**Fig. S2.**
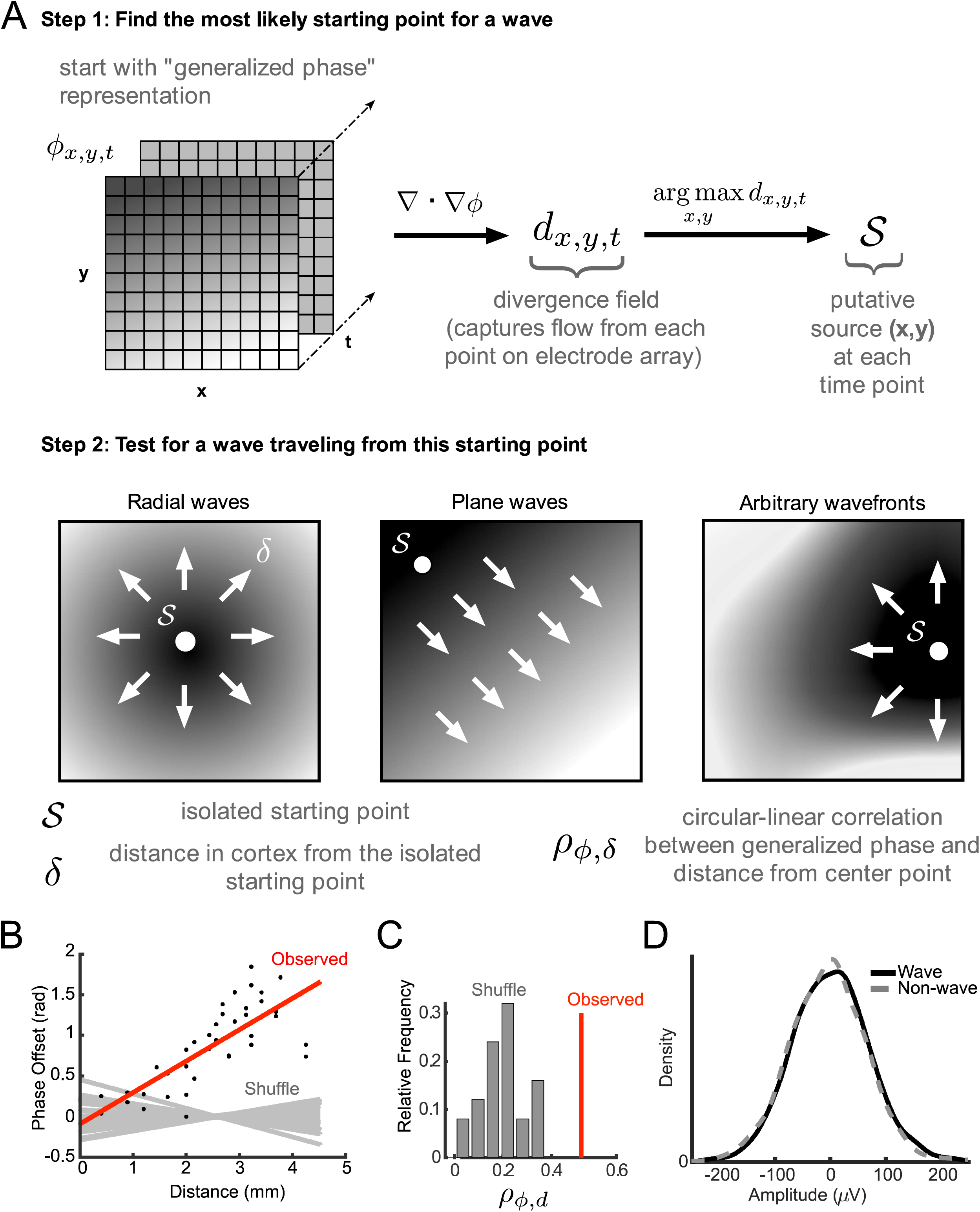
Detection of spontaneous traveling waves. **(A)** The method for detecting spontaneous waves from the generalized phase. First, the detection algorithm finds the most likely starting point for a putative wave as the point that maximizes the divergence of the phase gradient (step 1). With this source point found, the algorithm then quantifies the spatiotemporal organization about this point from the circular-linear correlation of phase with distance across the whole array (step 2). With this approach, the algorithm can robustly detect arbitrarily shaped wavefronts in the array data. **(B)** Scatter plot showing the phase offsets with distance from a putative source on the Utah array (black dots) and linear fit (red line). The linear fits for multiple instances of the same phase values randomly shuffled across the array have slopes much closer to zero (gray lines). **(C)** The distribution of shuffled phase correlations with distance (gray bars) determines the null hypothesis for which the observed putative wave (red line) must exceed to be classified as a wave. The critical phase correlation with distance value is determined by the 99^th^ percentile of the null distribution across all tested candidates. **(D)** There is no difference in the amplitude distribution of fluctuations that are detected as waves (black line), or rejected as waves (non-wave, gray dashed line).

**Fig. S3.**
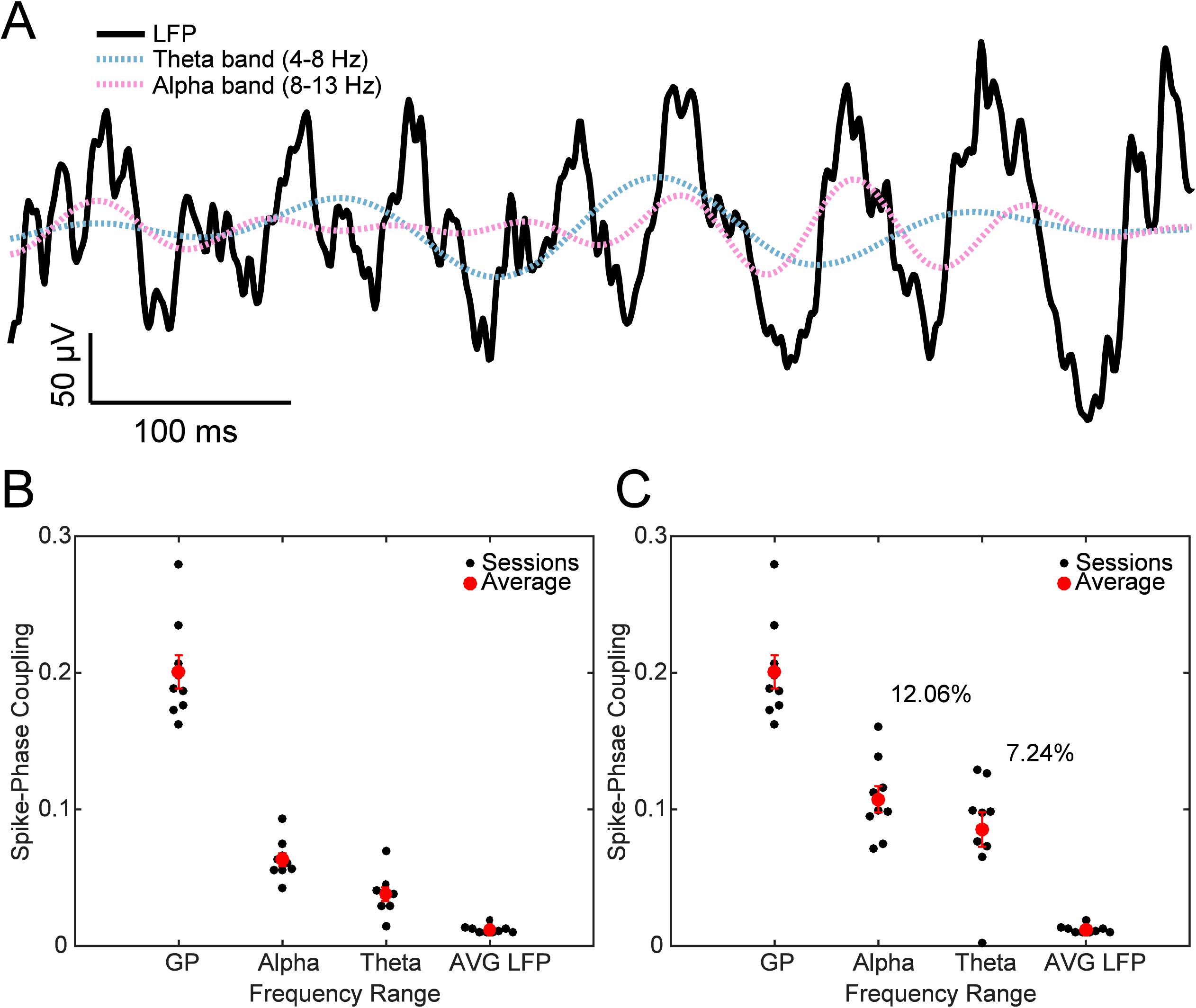
Wideband generalized phase is better coupled to spike timing than narrowband alpha or theta filters. **(A)** The phase and amplitude of the raw (5-100 Hz) LFP is poorly captured by narrow-band theta (4-8 Hz, blue dotted line) or alpha (8-13 Hz, red dotted line) filters. **(B)** Scatter plot showing spontaneous spike-phase coupling is greater for GP (5-40 Hz) than alpha or theta narrowband filtered phases. Average coupling across electrodes for Individual recording sessions are plotted as black dots and each red dot represents the average value across sessions. **(C)** Spontaneous spike-phase coupling remains stronger for GP than the narrow frequency bands even when the spontaneous LFP epochs are restricted to periods where there is large alpha (12.06% of recorded time) or theta (7.24%) LFP power during fixation (5 dB SNR, narrow-to broad-band power ratio). Results are presented from monkey W.

**Fig. S4.**
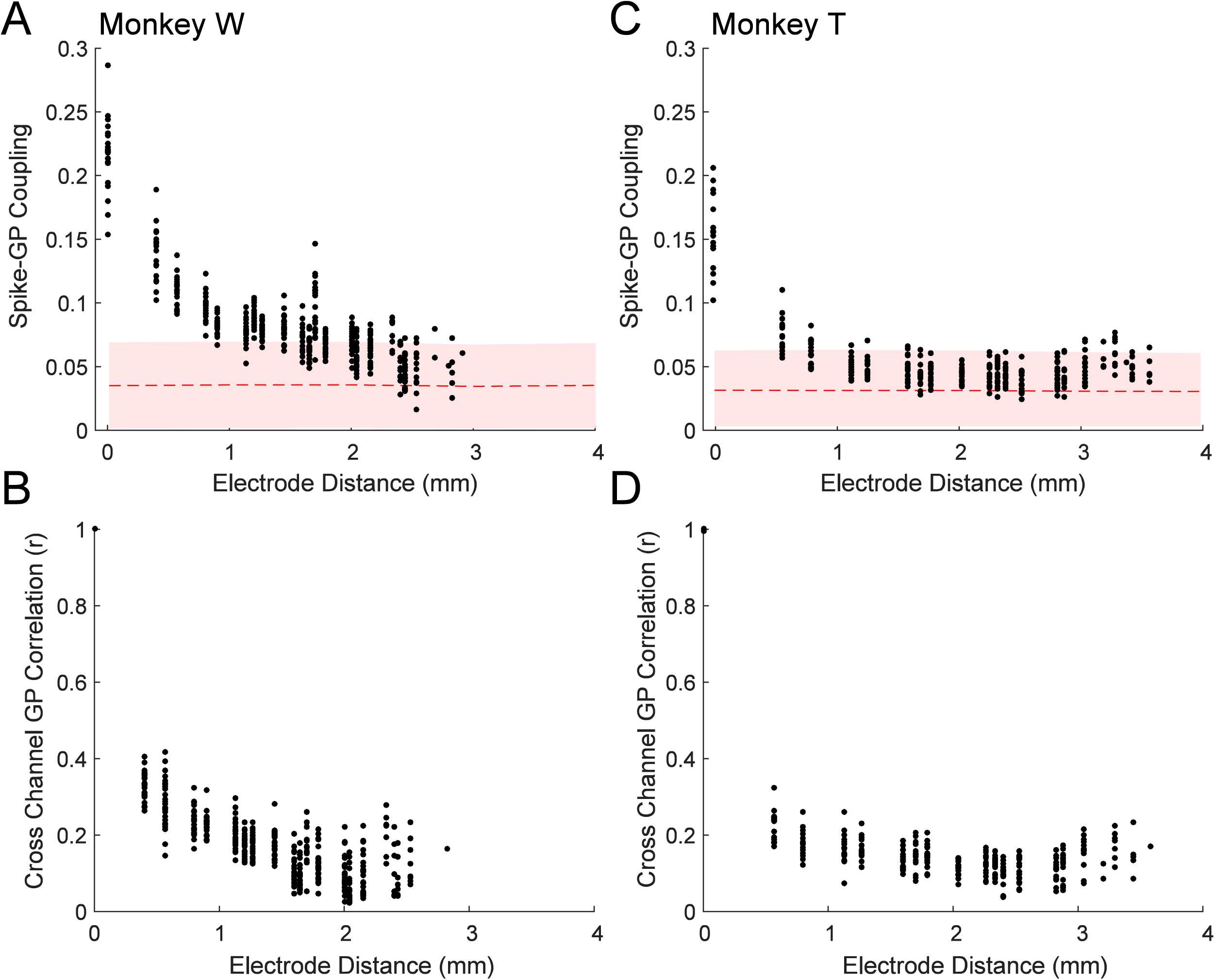
Spike coupling to generalized phase (GP) is spatially dependent. **(A)** Scatter plot showing the average spike-GP coupling across the distances of the array. Each point is averaged across a given spike-phase distance for a single recording session. The red dashed line shows the average null distribution for shuffled phases with 95 percent confidence interval (shaded region). Data shown for monkey W. **(B)** Scatter plot showing the cross-channel GP correlation for 100 ms of LFP during fixation across the electrode distances of the recording array. Each dot is the average correlation within an individual recording session at that channel distance. **(C,D)** Same as A and B, but for monkey T.

**Fig. S5.**
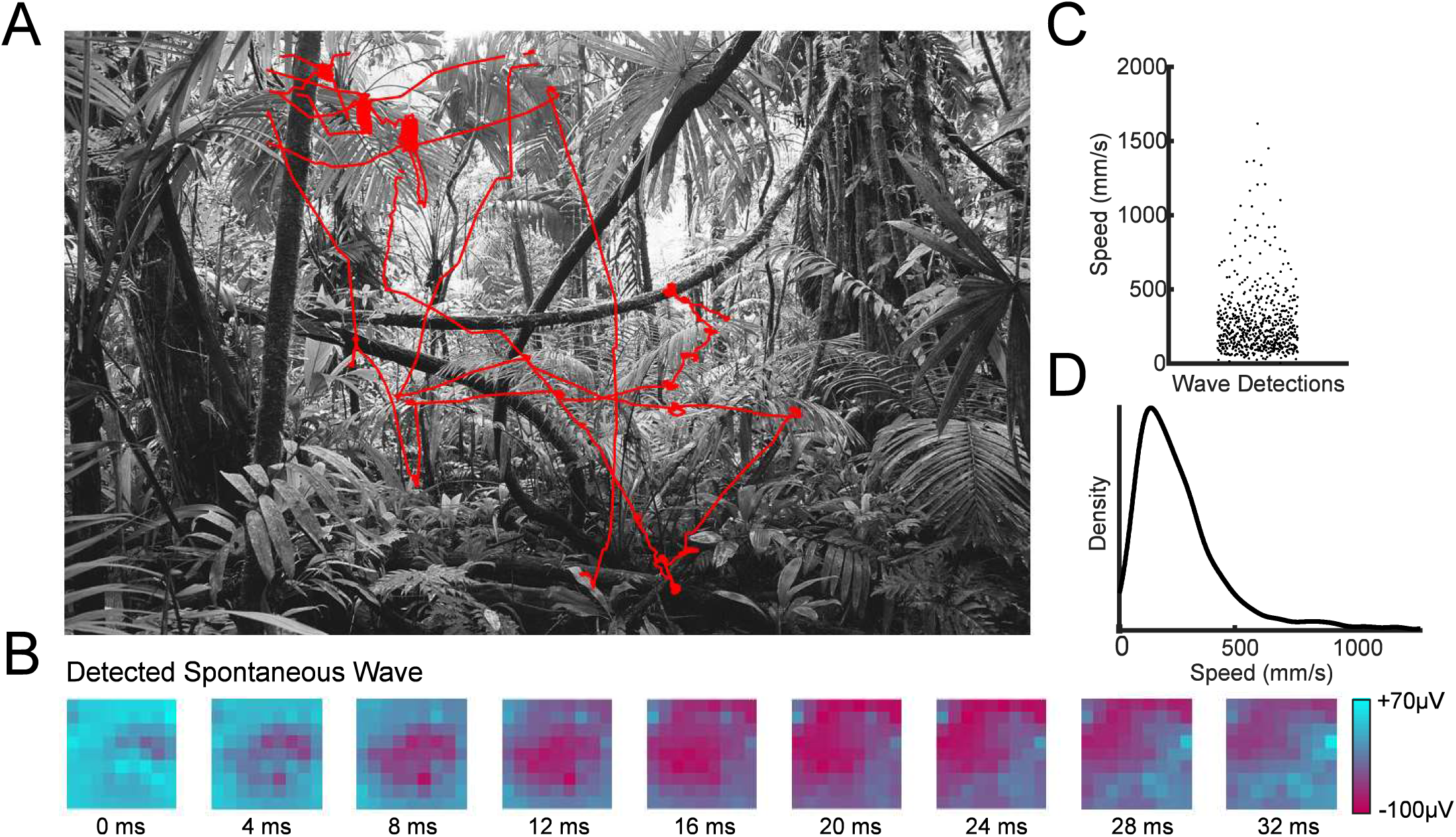
Spontaneous traveling waves are present during normal viewing of naturalistic visual scenes. Marmosets freely viewed static natural images for 10 seconds while head-fixed. **(A)** An example high contrast image with the gaze of the marmoset over the 10 second viewing interval shown in red. **(B)** An example of a spontaneous traveling wave detected during a period of fixation while monkey T was freely viewing a high contrast image. **(C)** Across 86 trials, 593 spontaneous traveling waves were detected during spontaneous fixations while the monkey freely viewed the images (−50 to +100 ms perisaccadic activity excluded). **(C)** The density of observed wave speeds was consistent with the conduction velocity of unmyelinated axons (100-600 mm/s).

**Fig. S6.**
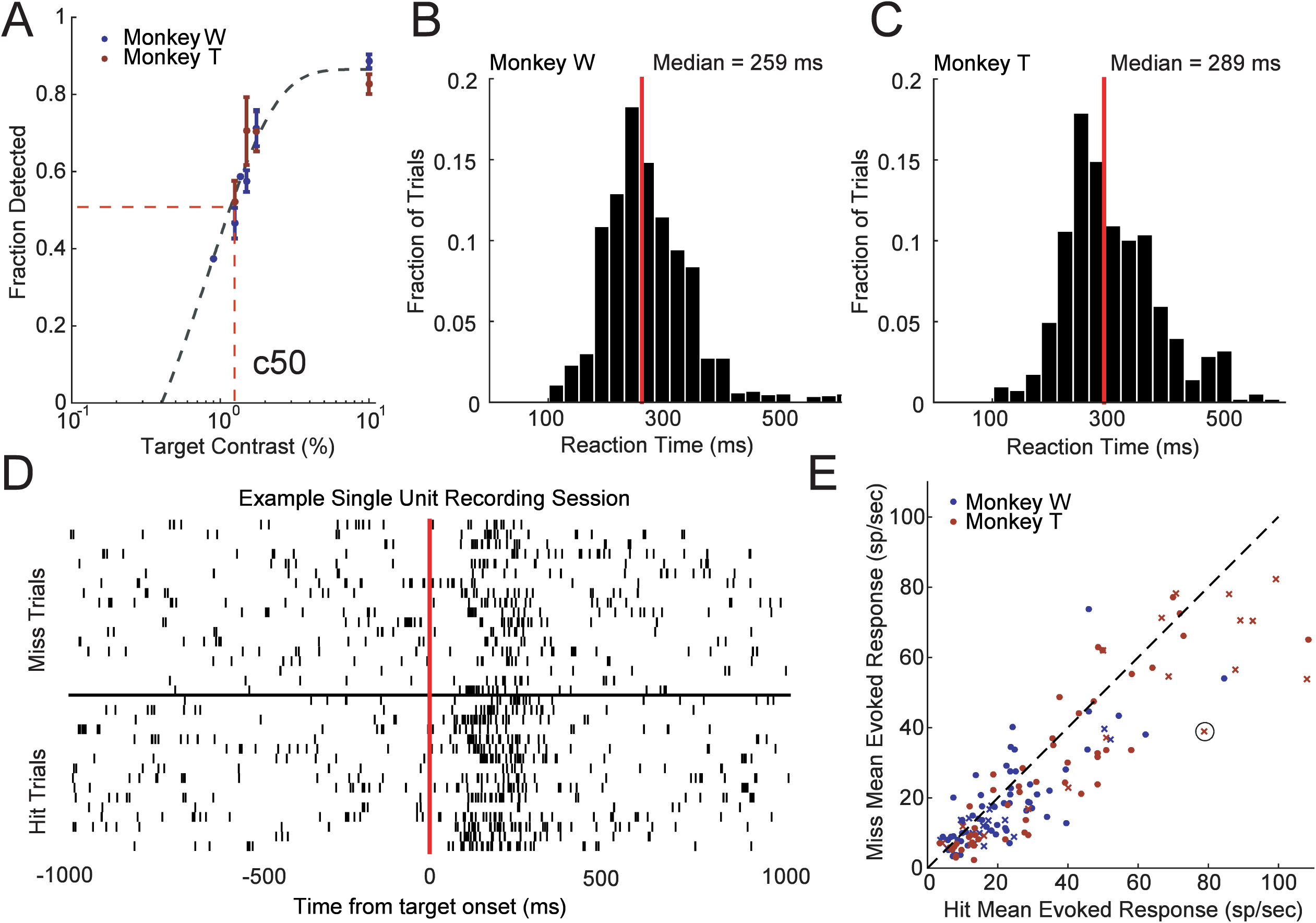
Target-evoked response magnitude is correlated with detection performance. **(A)** Detection performance of different target contrasts for monkey W (blue) and monkey T (red) across training days where those contrasts were presented. Both monkeys had similar psychophysical thresholds, defined as the contrast where the monkey detected the target 50 percent of the time on average (c50) as estimated from a sigmoid fit (gray dashed line). **(B, C)** Distributions of reaction times for monkey W (B) and monkey T (C) during the detection task at their c50 value. The median reaction time for each monkey is shown by a red line. **(D)** Spike rasters for an example neuron with trials sorted into hits (bottom rasters) and misses (top rasters). The target evoked firing rate was on average greater for hit trials than miss trials. **(E)** Scatter plot showing the distribution of mean hit (x-axis) and miss (y-axis) evoked responses (70-200 ms) for all single-(x’s) and multi-units (dots) recorded across all sessions for monkey W (blue) and monkey T (red). The circled x is the example neuron from panel D. Target-evoked responses were significantly stronger for detected targets in both monkeys.

**Fig. S7.**
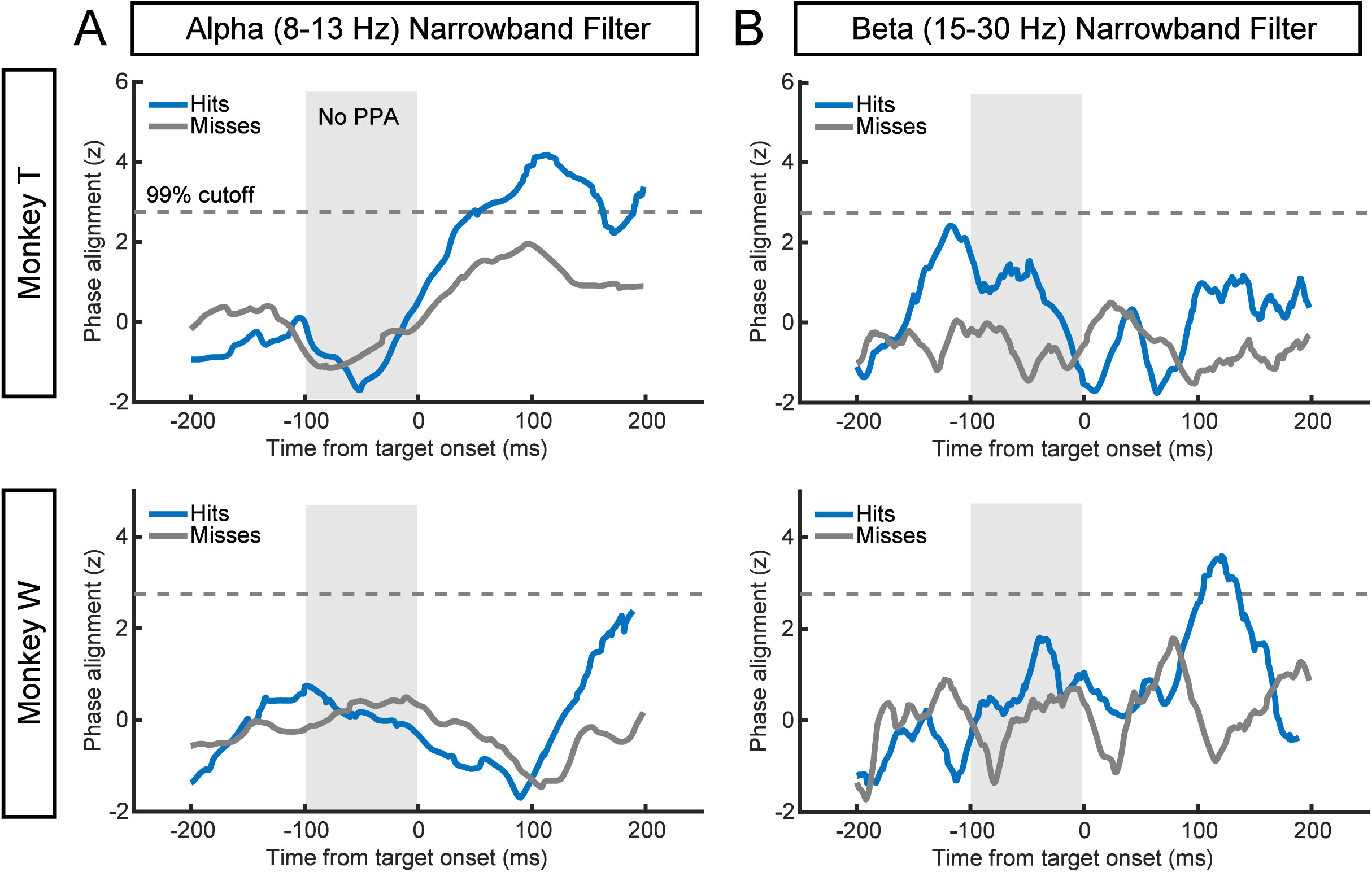
Narrowband filters fail to detect any significant wave phase alignment prior to target onset. **(A)** The cross-trial phase alignment computed as in Figure 3, but using a narrowband alpha (8-13 Hz) filter, does not show any significant alignment (gray dashed line) for hits (blue) or misses (gray) prior to target onset (gray region) for either monkey T (top) or monkey W (bottom). **(B)** The same as in (A), but for a beta (15-30 Hz) narrowband filter.

**Movie S1. Example of multiple detected traveling waves.** Video with 205 ms of spontaneous LFP data while monkey T is fixating a white dot on a gray screen. The wave examples in Figure 1B begin at 24 ms and 168 ms in this movie. Color axis scales from 80 µV (blue) to −80 µV (red).

## References

1. G. J. Tomko, D. R. Crapper, Neuronal variability: non-stationary responses to identical visual stimuli. Brain Res. 79, 405–418 (1974).

2. R. Vogels, W. Spileers, G. A. Orban, The response variability of striate cortical neurons in the behaving monkey. Exp. Brain Res. 77, 432–436 (1989).

3. M. N. Shadlen, W. T. Newsome, The variable discharge of cortical neurons: implications for connectivity, computation, and information coding. J. Neurosci. 18, 3870–3896 (1998).

4. M. Okun, A. Naim, I. Lampl, The subthreshold relation between cortical local field potential and neuronal firing unveiled by intracellular recordings in awake rats. J. Neurosci. 30, 4440–4448 (2010).

5. A. Luczak, P. Bartho, K. D. Harris, Gating of sensory input by spontaneous cortical activity. J. Neurosci. 33, 1684–1695 (2013).

6. A. Y. Y. Tan, Y. Chen, B. Scholl, E. Seidemann, N. J. Priebe, Sensory stimulation shifts visual cortex from synchronous to asynchronous states. Nature. 509, 226–229 (2014).

7. S. Katzner, I. Nauhaus, A. Benucci, V. Bonin, D. L. Ringach, M. Carandini, Local origin of field potentials in visual cortex. Neuron. 61, 35–41 (2009).

8. G. Buzsáki, C. A. Anastassiou, C. Koch, The origin of extracellular fields and currents—EEG, ECoG, LFP and spikes. Nat. Rev. Neurosci. 13, 407–420 (2012).

9. P. E. Roland, A. Hanazawa, C. Undeman, D. Eriksson, T. Tompa, H. Nakamura, S. Valentiniene, B. Ahmed, Cortical feedback depolarization waves: a mechanism of top-down influence on early visual areas. Proc. Natl. Acad. Sci. U. S. A. 103, 12586–12591 (2006).

10. W. Xu, X. Huang, K. Takagaki, J.-Y. Wu, Compression and reflection of visually evoked cortical waves. Neuron. 55, 119–129 (2007).

11. F. Han, N. Caporale, Y. Dan, Reverberation of recent visual experience in spontaneous cortical waves. Neuron. 60, 321–327 (2008).

12. I. Nauhaus, L. Busse, M. Carandini, D. L. Ringach, Stimulus contrast modulates functional connectivity in visual cortex. Nat. Neurosci. 12, 70–76 (2009).

13. A. Reimer, P. Hubka, A. K. Engel, A. Kral, Fast propagating waves within the rodent auditory cortex. Cereb. Cortex. 21, 166–177 (2011).

14. I. Ferezou, S. Bolea, C. C. H. Petersen, Visualizing the cortical representation of whisker touch: voltage-sensitive dye imaging in freely moving mice. Neuron. 50, 617–629 (2006).

15. S. Ray, J. H. R. Maunsell, Network rhythms influence the relationship between spike-triggered local field potential and functional connectivity. J. Neurosci. 31, 12674–12682 (2011).

16. D. Rubino, K. A. Robbins, N. G. Hatsopoulos, Propagating waves mediate information transfer in the motor cortex. Nat. Neurosci. 9, 1549–1557 (2006).

17. A. Benucci, R. A. Frazor, M. Carandini, Standing waves and traveling waves distinguish two circuits in visual cortex. Neuron. 55, 103–117 (2007).

18. L. Muller, A. Reynaud, F. Chavane, A. Destexhe, The stimulus-evoked population response in visual cortex of awake monkey is a propagating wave. Nat. Commun. 5, 3675 (2014).

19. T. P. Zanos, P. J. Mineault, K. T. Nasiotis, D. Guitton, C. C. Pack, A sensorimotor role for traveling waves in primate visual cortex. Neuron. 85, 615–627 (2015).

20. T. H. Bullock, M. C. Mcclune, J. T. Enright, Are the electroencephalograms mainly rhythmic? Assessment of periodicity in wide-band time series. Neuroscience. 121, 233–252 (2003).

21. H. Zhang, A. J. Watrous, A. Patel, J. Jacobs, Theta and Alpha Oscillations Are Traveling Waves in the Human Neocortex. Neuron. 98, 1269–1281.e4 (2018).

22. M. Steriade, A. Nuñez, F. Amzica, A novel slow (< 1 Hz) oscillation of neocortical neurons in vivo: depolarizing and hyperpolarizing components. J. Neurosci. 13, 3252–3265 (1993).

23. M. Steriade, I. Timofeev, F. Grenier, Natural waking and sleep states: a view from inside neocortical neurons. J. Neurophysiol. 85, 1969–1985 (2001).

24. L. Muller, G. Piantoni, D. Koller, S. S. Cash, E. Halgren, T. J. Sejnowski, Rotating waves during human sleep spindles organize global patterns of activity that repeat precisely through the night. Elife. 5 (2016), doi:10.7554/eLife.17267.

25. G. Buzsáki, A. Draguhn, Neuronal oscillations in cortical networks. Science. 304, 1926–1929 (2004).

26. C. Kayser, M. A. Montemurro, N. K. Logothetis, S. Panzeri, Spike-phase coding boosts and stabilizes information carried by spatial and temporal spike patterns. Neuron. 61, 597–608 (2009).

27. B. A. Lopour, A. Tavassoli, I. Fried, D. L. Ringach, Coding of information in the phase of local field potentials within human medial temporal lobe. Neuron. 79, 594–606 (2013).

28. M. J. McGinley, S. V. David, D. A. McCormick, Cortical Membrane Potential Signature of Optimal States for Sensory Signal Detection. Neuron. 87, 179–192 (2015).

29. T. P. Zanos, P. J. Mineault, C. C. Pack, Removal of spurious correlations between spikes and local field potentials. J. Neurophysiol. 105, 474–486 (2011).

30. D. Xing, C.-I. Yeh, R. M. Shapley, Spatial spread of the local field potential and its laminar variation in visual cortex. J. Neurosci. 29, 11540–11549 (2009).

31. V. Bringuier, F. Chavane, L. Glaeser, Y. Frégnac, Horizontal propagation of visual activity in the synaptic integration field of area 17 neurons. Science. 283, 695–699 (1999).

32. D. A. Lewis, Horizontal Synaptic Connections in Monkey Prefrontal Cortex. Cerebral Cortex Jan. 10, 82–92 (2000).

33. P. Girard, J. M. Hupé, J. Bullier, Feedforward and feedback connections between areas V1 and V2 of the monkey have similar rapid conduction velocities. J. Neurophysiol. 85, 1328–1331 (2001).

34. B. van Vugt, B. Dagnino, D. Vartak, H. Safaai, S. Panzeri, S. Dehaene, P. R. Roelfsema, The threshold for conscious report: Signal loss and response bias in visual and frontal cortex. Science. 360, 537–542 (2018).

35. C. C. H. Petersen, T. T. G. Hahn, M. Mehta, A. Grinvald, B. Sakmann, Interaction of sensory responses with spontaneous depolarization in layer 2/3 barrel cortex. Proc. Natl. Acad. Sci. U. S. A. 100, 13638–13643 (2003).

36. J. F. A. Poulet, C. C. H. Petersen, Internal brain state regulates membrane potential synchrony in barrel cortex of behaving mice. Nature. 454, 881–885 (2008).

37. N. A. Busch, J. Dubois, R. VanRullen, The phase of ongoing EEG oscillations predicts visual perception. J. Neurosci. 29, 7869–7876 (2009).

38. K. E. Mathewson, G. Gratton, M. Fabiani, D. M. Beck, T. Ro, To See or Not to See: Prestimulus α Phase Predicts Visual Awareness. J. Neurosci. 29, 2725–2732 (2009).

39. J. Samaha, O. Gosseries, B. R. Postle, Distinct Oscillatory Frequencies Underlie Excitability of Human Occipital and Parietal Cortex. J. Neurosci. 37, 2824–2833 (2017).

40. I. C. Fiebelkorn, Y. B. Saalmann, S. Kastner, Rhythmic sampling within and between objects despite sustained attention at a cued location. Curr. Biol. 23, 2553–2558 (2013).

41. R. F. Helfrich, I. C. Fiebelkorn, S. M. Szczepanski, J. J. Lin, J. Parvizi, R. T. Knight, S. Kastner, Neural Mechanisms of Sustained Attention Are Rhythmic. Neuron. 99, 854–865.e5 (2018).

42. I. C. Fiebelkorn, M. A. Pinsk, S. Kastner, A Dynamic Interplay within the Frontoparietal Network Underlies Rhythmic Spatial Attention. Neuron. 99, 842–853.e8 (2018).

43. M. G. P. Rosa, G. N. Elston, Visuotopic organisation and neuronal response selectivity for direction of motion in visual areas of the caudal temporal lobe of the marmoset monkey (Callithrix jacchus): middle temporal area, middle temporal crescent, and surrounding cortex. J. Comp. Neurol. 393, 505–527 (1998).

44. D. Gabor, Theory of communication. Part 1: The analysis of information. Journal of the Institution of Electrical Engineers - Part III: Radio and Communication Engineering. 93, 429–441 (1946).

45. M. Le Van Quyen, J. Foucher, J. Lachaux, E. Rodriguez, A. Lutz, J. Martinerie, F. J. Varela, Comparison of Hilbert transform and wavelet methods for the analysis of neuronal synchrony. J. Neurosci. Methods. 111, 83–98 (2001).

46. A. V. Oppenheim, R. W. Schafer, J. R. Buck, Discrete-time Signal Processing (Prentice Hall, 1999).

47. L. Marple, Computing the discrete-time“ analytic” signal via FFT. IEEE Trans. Signal Process. 47, 2600–2603 (1999).

48. M. Johansson, The hilbert transform. Mathematics Master’s Thesis. Växjö University, Suecia. Disponible en internet: http://w3.msi.vxu.se/exarb/mj_ex.pdf, consultado el. 19 (1999) (available at http://www.fuchs-braun.com/media/d9140c7b3d5004fbffff8007fffffff0.pdf).

49. M. Feldman, Hilbert transform in vibration analysis. Mech. Syst. Signal Process. 25, 735–802 (2011/4).

50. E. Pereda, A. Gamundi, R. Rial, J. González, Non-linear behaviour of human EEG: fractal exponent versus correlation dimension in awake and sleep stages. Neurosci. Lett. 250, 91–94 (1998).

51. K. Linkenkaer-Hansen, V. V. Nikouline, J. M. Palva, R. J. Ilmoniemi, Long-range temporal correlations and scaling behavior in human brain oscillations. J. Neurosci. 21, 1370–1377 (2001).

52. J. Milstein, F. Mormann, I. Fried, C. Koch, Neuronal shot noise and Brownian 1/f2 behavior in the local field potential. PLoS One. 4, e4338 (2009).

53. S. Rao Jammalamadaka, A. Sengupta, Topics in Circular Statistics (World Scientific, 2001).

54. T. D. Frank, M. J. Richardson, On a test statistic for the Kuramoto order parameter of synchronization: An illustration for group synchronization during rocking chairs. Physica D. 239, 2084–2092 (2010).

